# The enzyme Rnf completes the pathway for forming propionate during fermentation in *Prevotella*

**DOI:** 10.1101/2022.11.08.515646

**Authors:** Bo Zhang, Christopher Lingga, Hannah De Groot, Timothy J. Hackmann

**Affiliations:** Department of Animal Science, University of California, Davis, CA, USA

## Abstract

Propionate is a microbial metabolite that is formed in the gastrointestinal tract, and it affects host physiology as a source of energy and signaling molecule. Despite the importance of propionate, the biochemical pathways responsible for its formation are not clear in all microbes. For the succinate pathway used during fermentation, a key enzyme appears to be missing—one that can oxidize ferredoxin and reduce NAD. Here we show that Rnf [ferredoxin--NAD(+) oxidoreductase (Na(+)-transporting)] is this key enzyme in two abundant bacteria of the rumen (*Prevotella brevis* and *Prevotella ruminicola*). We found these bacteria form propionate, succinate, and acetate with the classic succinate pathway. At first, this pathway appears unbalanced, forming reduced ferredoxin and oxidized NAD in excess. If this continued unabated, fermentation would halt within 1.5 s. We found these bacteria solve this problem by oxidizing ferredoxin and reducing NAD with Rnf. This is demonstrated using growth experiments, genomics, proteomics, and enzyme assays. Genomic and phenotypic data suggest many bacteria use Rnf similarly. We cataloged fermentation products of >1,400 species of prokaryotes, and nearly 10% formed propionate, succinate, and acetate. Over 40% of species carrying out this fermentation also had genes for Rnf. This work shows Rnf is important to propionate formation in many bacteria from the environment, and it provides fundamental knowledge for manipulating fermentative propionate production.

## INTRODUCTION

Metabolites formed via anaerobic fermentation in the gastrointestinal of mammals have great effects on the host physiology and health (1, 2). The major metabolites formed by gut bacteria during fermentation of dietary carbohydrates are short-chain fatty acids (SCFAs). As one of the major SCFAs, propionate can affect satiety and glucose homeostasis in humans (3, 4). It has beneficial effects on beta-cell function to maintain healthy glucose homeostasis (5). Recently, it has been demonstrated that propionate can suppress colorectal cancer growth (6, 7), while excess levels of propionate may lead to Alzheimer’s disease by inducing hyperammonemia (8). Furthermore, propionate also plays important roles in other animals, such as ruminants. It is a major source of glucose for the ruminants, and about 50% of glucose is from propionate (9). Propionate formation in the rumen of ruminants is negatively related with methane emission, since they compete for metabolic hydrogen in the rumen. Favoring propionate formation could mitigate methane emissions (10). Realizing its importance in human health, agricultural production, and the environment, studies focusing on biochemical pathways have revealed many enzymes responsible for fermentative propionate production (11, 12).

Three biochemical pathways are responsible for fermentative propionate production from dietary carbohydrates, including the succinate pathway, the acrylate pathway, and the propanediol pathway (11, 12). Propionate is most commonly formed using the succinate pathway and in combination with acetate. This pathway involves the conversion of succinate to propionate via methylmalonyl-CoA. Some organisms with this pathway include *Bacteroides fragilis* (13) and *Selenomonas ruminantium* (14). The acrylate pathway involves the conversion of lactate to lactoyl-CoA, acryloyl-CoA, propionyl-CoA and propionate, e.g. *Coprococcus catus* (11) and *Megasphaera elsdenii* (15). Some gut commensal bacteria, such as *Roseburia inulinivorans* (16), carry out the propanediol pathway to from propionate from deoxy sugars.

Despite decades of study, one major biochemical pathway (i.e. the succinate pathway) for forming propionate has unknown steps. When glucose is the substrate, this pathway has a problem: it forms excess amounts of reduced ferredoxin (Fd_red_), a redox cofactor (Fig. 1A) (17). This cofactor is formed by the enzyme pyruvate:ferredoxin oxidoreductase (EC 1.2.7.1), and no step is known to oxidize it back to ferredoxin (Fd_ox_) during this pathway. Similarly, the pathway forms oxidized NAD (NAD_ox_), with no step to reduce it back to reduced NAD (NAD_red_). This is an apparent problem in both prokaryotes (17, 18) and eukaryotes (19). These unknown steps are important because if Fd_ox_ and NAD_red_ is exhausted (not regenerated), fermentation will halt.

**Fig. 1.**
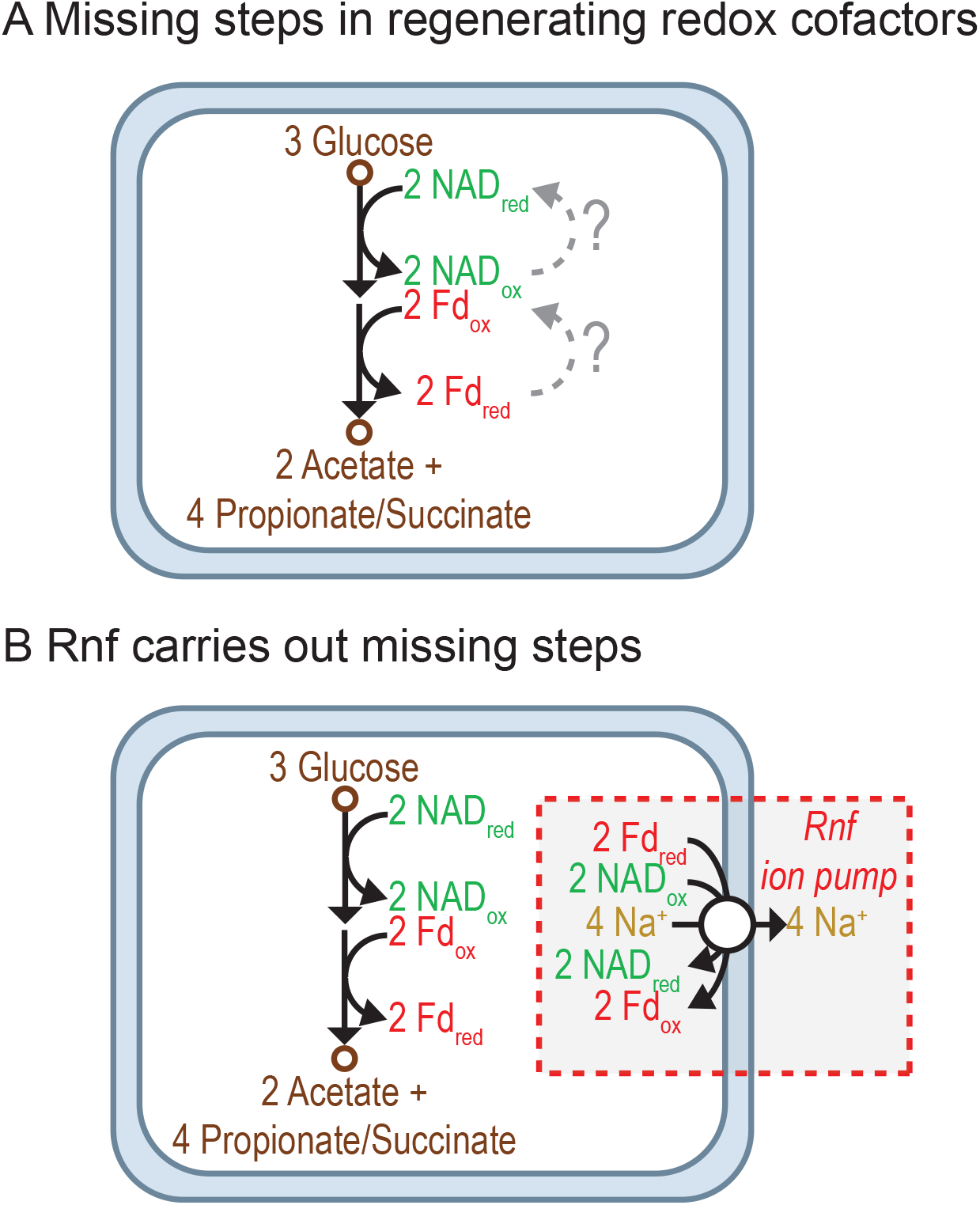
Fermentation of glucose to propionate, succinate, and acetate has missing or unknown steps. (A) The missing steps are for regenerating redox cofactors. (B) We hypothesize Rnf carries out the missing steps. Abbreviations: Fd_ox_, oxidized ferredoxin; Fd_red_, reduced ferredoxin (two reduced iron-sulfur clusters); NAD_ox_, oxidized NAD; NAD_red_, reduced NAD.

We hypothesized that the enzyme Rnf fills in the missing steps (Fig. 1B). The enzyme Rnf [ferredoxin--NAD(+) oxidoreductase (Na(+)-transporting), EC 7.2.1.2] simultaneously oxidizes Fd_red_ and reduces NAD_ox_, solving two problems at once. This enzyme plays a similar role in other pathways, such as one metabolizing caffeate (20). Recently, we found Rnf genes in many propionate-forming bacteria from the rumen (17). Here we study two of these rumen bacteria in detail and find that they indeed use Rnf in forming propionate (or its precursor, succinate). We show this using growth experiments, genomics, proteomics, and enzyme assays. Further, we find Rnf is common in bacteria that form propionate (or succinate), with 44 type strains from many habitats encoding it. This work suggests Rnf is important to propionate formation in many bacteria from the environment.

## RESULTS

### *Prevotella* form propionate, succinate, and acetate during fermentation

Our hypothesis was that fermentation of glucose to propionate, succinate, and acetate uses Rnf. We studied this in two bacteria from the rumen (*Prevotella brevis* GA33 and *Prevotella ruminicola* 23). The first step was to verify that these organisms form propionate, succinate, and acetate and in the ratios expected (Fig. 1). We grew these bacteria on media containing glucose and ammonia, then analyzed the culture for several products. For *P. brevis* GA33, we used a medium that also contained yeast extract and trypticase, as it would not grow on media with glucose only.

Both species formed large amounts of succinate and acetate (Fig. 2A). Propionate was formed in large amounts by *P. ruminicola* 23, whereas it was formed in only trace amounts by *P. brevis* GA33 (Fig. S1). The ratio of succinate and propionate to acetate was approximately 2:1. They also formed formate, D-lactate, and L-lactate, but only in trace amounts. These results follow our expectations (Fig. 1).

**Fig. 2.**
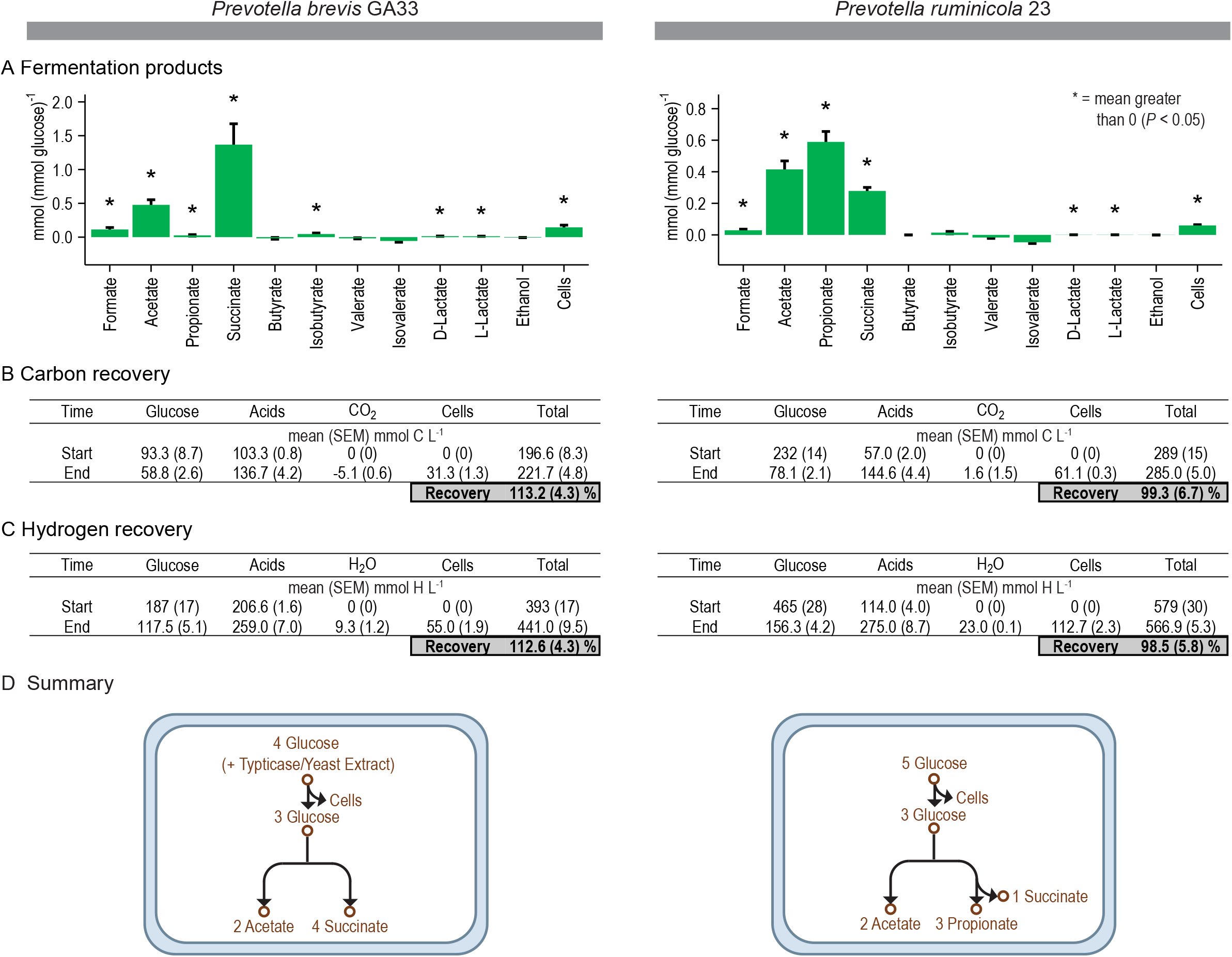
*Prevotella* form propionate, succinate, and acetate during fermentation of glucose. (A) Yield of fermentation products. (B) Recovery of carbon is near or above 100%. (C) Recovery of hydrogen is also near or above 100%. (D) Summary of growth and fermentation. In (A), the yield of cells is g (mmol glucose)^-1^. Results are mean ± standard error of at least 3 biological replicates (culture supernatant or cells prepared from independent cultures).

Neither species formed H_2_ (Fig. S2). As a control, we analyzed gas samples from *S. ruminantium* HD4, a propionate-forming bacterium that forms H_2_ in trace amounts (21). We were indeed able to detect H_2_ formation by this organism (Fig. S2). This result shows that if *Prevotella* formed H_2_, even in trace amounts, we would have been able to detect it.

To check how accurately we measured products, we calculated carbon and hydrogen recovery (Fig. 2B and C). A recovery of 100% indicates that all carbon (or hydrogen) at the start of incubation was recovered in products measured at the end. We found recoveries were near or above 100%. For *P. brevis* GA33, values were above 100% because our calculations did not account for trypticase and yeast extract also in the medium of this bacterium. The high recoveries of carbon and hydrogen indicate that we measured all products accurately.

In sum, our work shows that *Prevotella* form propionate, succinate, and acetate as the sole products of fermentation (Fig. 2D). Additionally, they form these products in the ratio expected (Fig. 1).

### Fermentation in *Prevotella* appears to be unbalanced

We hypothesized that *Prevotella* use Rnf to balance fermentation. Without this enzyme, fermentation should produce excess NAD_ox_ and Fd_red_. We determined if this was indeed the case for *Prevotella*.

We calculated the quantity of NAD_ox_ and Fd_red_ produced during the experiments above (Table S1). Our calculations revealed that excess NAD_ox_ and Fd_red_ were indeed formed. The amount was 2.7 NAD_ox_ and 2.3 Fd_red_ per 3 glucose (Table S1). This was even higher than expected (Fig. 1) and owed to additional NAD_ox_ and Fd_red_ being formed during production of cells (particularly lipid) (Table S2). This calculation did not include the activity of Rnf, and it shows without this enzyme, fermentation would indeed be unbalanced.

Next, we calculated how long cells could sustain such an unbalanced fermentation. We calculated that all Fd_ox_ would be consumed and fermentation would halt within 1.5 s. This calculation assumes 2.3 Fd_red_ per 3 glucose fermented (Table S1), 667 nmol glucose fermented (g dry cells)^-1^ s^-1^ (see Materials and Methods), 75 nmol total ferredoxin/g wet cells (22), wet cells are 10% dry mass, and all ferredoxin starts as Fd_ox_. Without Rnf, cells could sustain fermentation only for seconds (or less).

We performed calculations on *P. ruminicola* 23 only. To calculate the quantity of NAD_ox_ and Fd_red_ formed during production of cells, we assumed macromolecules were synthesized from glucose and ammonia (Table S2). This would have been a bad assumption for *P. brevis* GA33, where macromolecules could have come from trypticase and yeast extract.

Our calculation points to an apparent excess of NAD_ox_ and Fd_red_ formed during fermentation and growth. It shows a critical need for Rnf or a similar enzyme.

### *Prevotella* have Rnf [ferredoxin--NAD(+) oxidoreductase (Na(+)-transporting)]

Having established a need for an enzyme like Rnf, we determined if this activity of Rnf is indeed possessed by *Prevotella*. Genomics, proteomics, and enzyme assays were used to test its presence.

We found Rnf in both the genome and proteome (Fig. 3, Table S3, Table S4). The genomes of both species had genes for all six subunits of this enzyme (Fig. 3A). Proteomics revealed genes for four subunits were expressed in *P. brevis* GA33 and three in *P. ruminicola* 23 (Fig. 3B). Our methods were exhaustive and used multiple sample types (cell extract, cell membrane) and acquisition methods (data-dependent acquisition, data-independent acquisition). The two subunits we never detected (RnfA and RnfE) are predicted to be integral proteins, which are challenging targets in proteomics. These subunits have evaded detection even in purified Rnf (23).

**Fig. 3.**
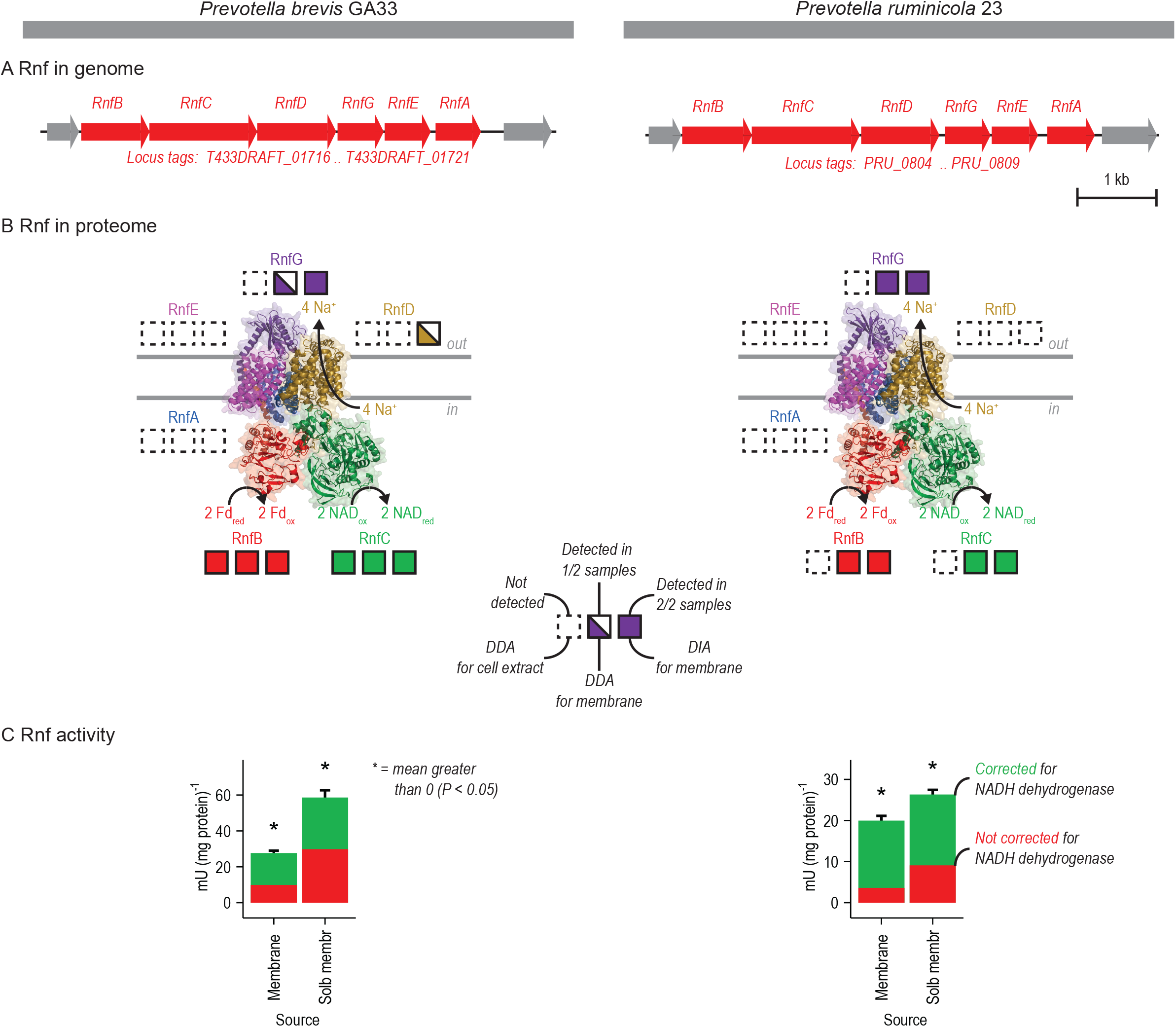
*Prevotella* have the enzyme Rnf. Rnf is evident in the (A) genome, (B) proteome, and (C) measurements of enzyme activity. Results in (C) are mean ± standard error of 3 biological replicates (cell membranes prepared from independent cultures). To correct Rnf for activity of NADH dehydrogenase, we added the activity measured for Nqr (see Table 2). Abbreviations: Fd_ox_, oxidized ferredoxin; Fd_red_, reduced ferredoxin (two reduced iron-sulfur clusters); NAD_ox_, oxidized NAD; NAD_red_, reduced NAD; DDA, data-dependent acquisition; DIA, data-independent acquisition; Membrane, cell membrane sample; Solb membr, solubilized cell membrane sample. See Tables S3 and S4 for more information.

After finding evidence of Rnf in the genome and proteome, we tested for its catalytic activity [ferredoxin--NAD(+) oxidoreductase] with enzyme assays (Fig. 3C). To do so, we measured formation of NAD_red_ by cell membranes after adding Fd_red_. Using these assays, we found that cell membrane of both *Prevotella* species had activity (Fig. 3C). The activity depended on adding both Fd_red_ and NAD_ox_. Further, activity was localized to the membrane; activity in cytoplasmic contents was low for *P. brevis* GA33 (3.1 [0.5] (mean [standard error of mean]) mU/mg) and undetectable for *P. ruminicola* 23. These experiments show *Prevotella* had activity of Rnf, and the properties were as expected. Likewise, these experiments rule out the presence of a similar enzyme in the cytoplasm [a cytoplasmic ferredoxin--NAD(+) oxidoreductase].

At first, activity observed in the cell membrane appeared low. However, we found higher activity after correcting for activity of NADH dehydrogenase that leads to consumption of NAD_red_ formed in the assay (Fig. 3C). This correction is commonly done for other NADH-dependent enzymes [see, for example, Asanuma and Hino (24)], and it makes sense to do it with Rnf. We found still higher activity after performing a partial purification of Rnf (by solubilizing cell membranes in detergent) (Fig. 3C). This shows that activity, at first low, is indeed present and on par with other membrane-bound enzymes (see below).

To verify that this activity was due to Rnf, not another enzyme, we determined its dependence on sodium ion (Fig. 4). In most species, Rnf pumps sodium ions (to create a gradient) and thus depends on them for high activity (23, 25, 26). We found that *P. brevis* GA33 did not grow without sodium, showing a general dependence on this ion (Fig. 4A). We found the same for *P. ruminicola* 23 (data not shown). Further, when we directly tested if sodium ion stimulated the catalytic activity of Rnf, we found that it did (Fig. 4B).

**Fig. 4.**
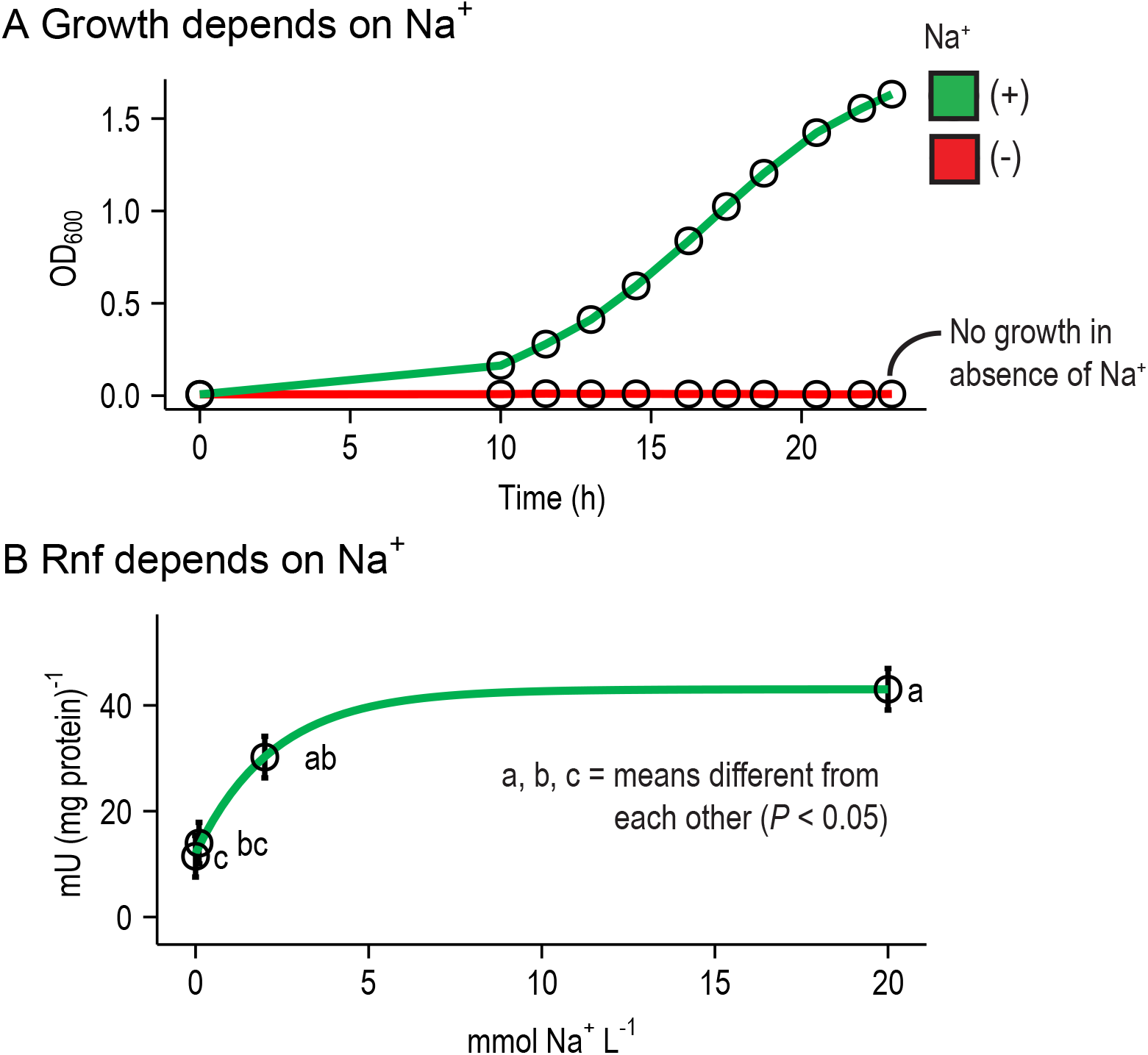
*Prevotella brevis* GA33 depends on Na^+^ for growth and for Rnf activity. In (A), sodium was removed from the media by replacing NaCl, NaOH and Na_2_CO_3_ with equimolar KCl, KOH, and K_2_CO_3_. Resazurin was also removed. Results are for one representative culture. Experiments were done with 2 cultures on 2 different days. In (B), sodium was removed from the assay mix by replacing NAD sodium salt, sodium pyruvate, and CoA lithium salt with equimolar NAD hydrate, potassium pyruvate, and CoA hydrate. The residual Na^+^ in the Tris-Cl buffer and MgCl_2_ was 2 μM (as measured by an electrode; Fisher Accumet 13-620-503A). No correction was made for NADH dehydrogenase activity. Results are mean ± standard error of 4 biological replicates (cell membranes prepared from independent cultures).

Our enzyme assays required Fd_red_, which we generated using a system similar to Schoelmerich et al. (27). Specifically, we purified ferredoxin from *C. pasteurianum* 5, then we reduced it with pyruvate and crude pyruvate:ferredoxin oxidoreductase. The crude pyruvate:ferredoxin oxidoreductase was cytoplasmic contents from the organism in which Rnf was tested (*P. brevis* GA33 or *P. ruminicola* 23). We verified that the crude pyruvate:ferredoxin oxidoreductase worked as intended. First, we used it to detect activity of Rnf in *Pseudobutyrivibrio ruminis* A12-1. We found activity of 50.0 [1.9] mU/mg, which is similar to the value found by schoelmerich et al. (27) for the same organism. Second, we screened it for activity of interfering enzymes, including a cytoplasmic ferredoxin--NAD(+) oxidoreductase and pyruvate dehydrogenase. We found these interfering activities were low or undetectable (see results above for cytoplasmic ferredoxin--NAD(+) oxidoreductase and see below for pyruvate dehydrogenase). These results show this system is appropriate for generating Fd_red_ in Rnf assays.

In sum, our work at the genomic proteomic, and enzymatic level establishes that *Prevotella* have Rnf. With it, *Prevotella* can handle excess NAD_ox_ and Fd_red_ produced during fermentation.

### *Prevotella* have other enzymes needed to form fermentation products

After finding that Rnf was present in *Prevotella*, we determined if other enzymes forming propionate, succinate, and acetate were also present. This was important to confirm that redox cofactors (NAD_ox_ and Fd_red_) are produced in the pathway as expected. Again, we relied on genomics, proteomics, and enzyme assays.

We found enzymes of the classic succinate pathway in the genome and proteome (Fig. 5, Table S3, Table S4). When using proteomics, we found cytoplasmic enzymes were well detected (Fig. 5A). Membrane-bound proteins were also detected, though some subunits (corresponding to integral proteins) were missed (as with Rnf) (Fig. 5B). The membrane-bound proteins Nqr (EC 7.2.1.1) and fumarate reductase (EC 1.3.5.1) in our bacteria have also been detected in *Prevotella bryantii* B_1_4 (28), where they have been characterized. Together, these enzymes form a pathway where Rnf is needed to regenerate NAD_red_ and Fd_ox_.

**Fig. 5.**
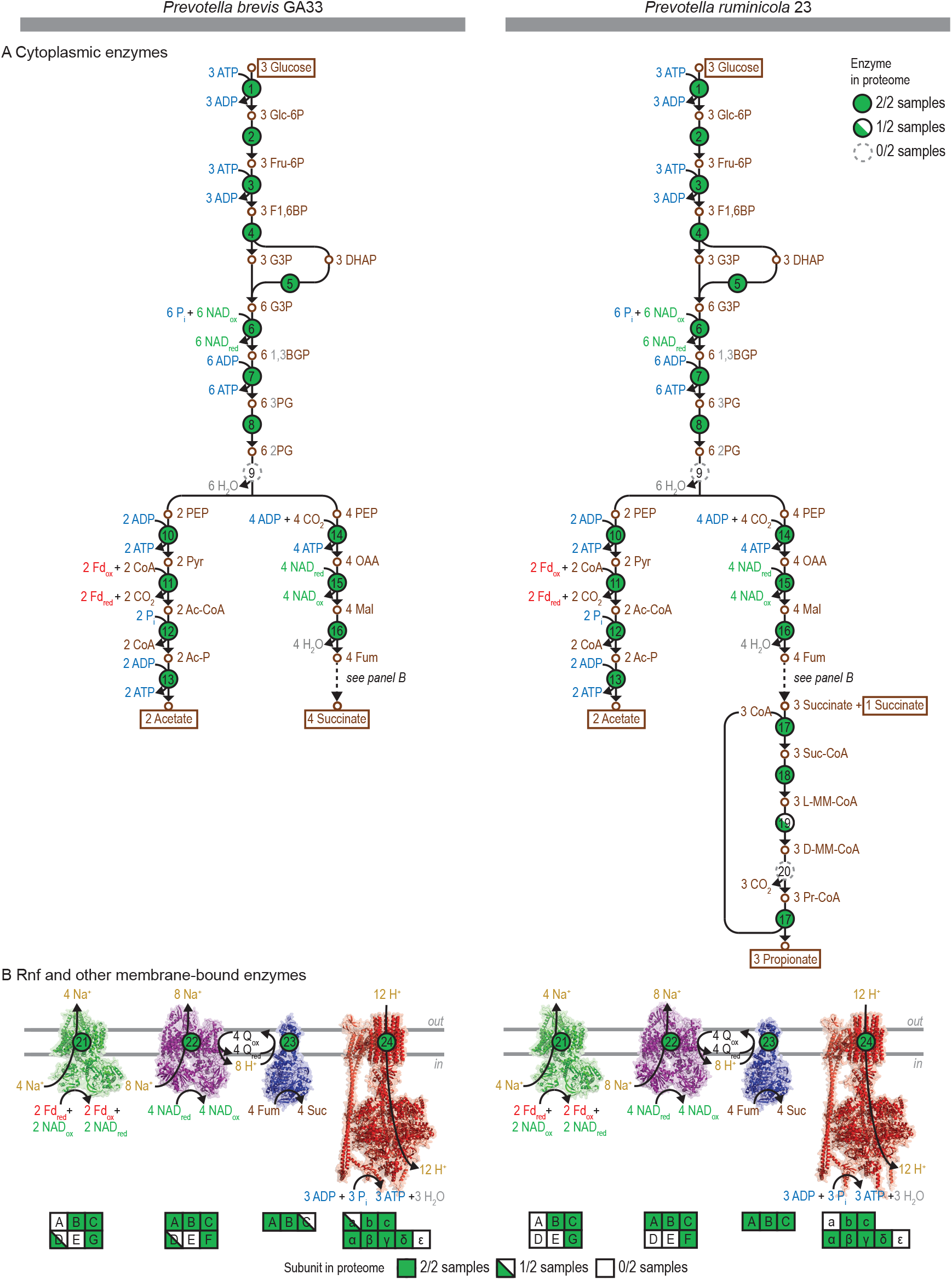
*Prevotella* have enzymes for forming propionate, succinate, and acetate in the proteome. (A) Cytoplasmic enzymes. (B) Rnf and other membrane-bound enzymes. Abbreviations: Glc-6P, glucose-6-phosphate; Fru-6P, fructose-6-phosphate; F1,6BP, fructose-1,6-bisphophate; G3P, glyceraldehyde-3-phosphate; DHAP, dihydroxyacetone phosphate; 1,3BGP, 1,3-bisphosphoglycerate; 3PG, 3-phosphoglycerate; 2PG, 2-phosphoglycerate; PEP, phosphoenolpyruvate; Pyr, pyruvate; Ac-CoA, acetyl-CoA; Ac-P, acetyl-phosphate; OAA, oxaloacetate; Mal, malate; Fum, fumarate; Suc-CoA, succinyl-CoA; L-MM-CoA, L-methylmalonyl-CoA; D-MM-CoA, D-methylmalonyl-CoA; Pr-CoA, propionyl-CoA; Fd_ox_, oxidized ferredoxin; Fd_red_, reduced ferredoxin (two reduced iron-sulfur clusters); NAD_ox_, oxidized NAD; NAD_red_, reduced NAD; CoA, coenzyme A; P_i_, inorganic phosphate; Q_ox_, oxidized quinone; Q_red_, reduced quinone. See Tables S3 and S4 for more information.

There was one enzyme missing in *P. brevis* GA33 and another in *P. ruminicola* 23 (Fig. 5A). In *P. brevis* GA33, an enzyme of glycolysis (enolase) was missing in the genome and proteome. This has no easy explanation but has been found previously in the genome (17, 29). In *P. ruminicola* 23, an enzyme for converting succinate to propionate was likewise missing. Despite this finding, there is evidence the enzyme is present (or substituted by a similar enzyme). The conversion of succinate to propionate is well known to require vitamin B_12_ (30), and *P. ruminicola* 23 formed propionate only when this vitamin was in the media (data not shown). Others have found the same for this bacterium (31). With a few possible exceptions, our work shows that *Prevotella* have the expected enzymes in the genome and proteome.

After finding evidence in the genome and proteome, we tested for catalytic activity of key enzymes (Table 2). We focused mostly on enzymes that generate redox cofactors used by Rnf. We detected activity in all cases expected. For example, we found activity of malate dehydrogenase (EC 1.1.1.37), which produces NAD_ox_ (used by Rnf). A similar enzyme producing NADP_ox_ (EC 1.1.1.82) was also detected, but with lower activity. Our work confirms that *Prevotella* have the expected enzymes for forming propionate, succinate, and acetate—including those that form NAD_ox_ and Fd_red_. Rnf is needed to complete this pathway.

### Rnf is important in many organisms forming propionate, succinate, and acetate

We wanted to see if Rnf is distributed widely in organisms that form propionate, succinate, and acetate. To do so, we used genomic and phenotypic data for prokaryotes from *Bergey’s Manual of Systematics of Archaea and Bacteria* (Table S5).

We first determined how many prokaryotes form propionate, succinate, and acetate during fermentation. We constructed a heat map to summarize fermentation products reported for organisms in *Bergey’s Manual* (Fig. 6). This encompasses 39 products from over 1,400 type strains (Table S5). We found that prokaryotes that form exclusively propionate/succinate and acetate (no other products) represent about 10% of the total. Thus, fermentations that form propionate, succinate, and acetate are common.

**Fig. 6.**
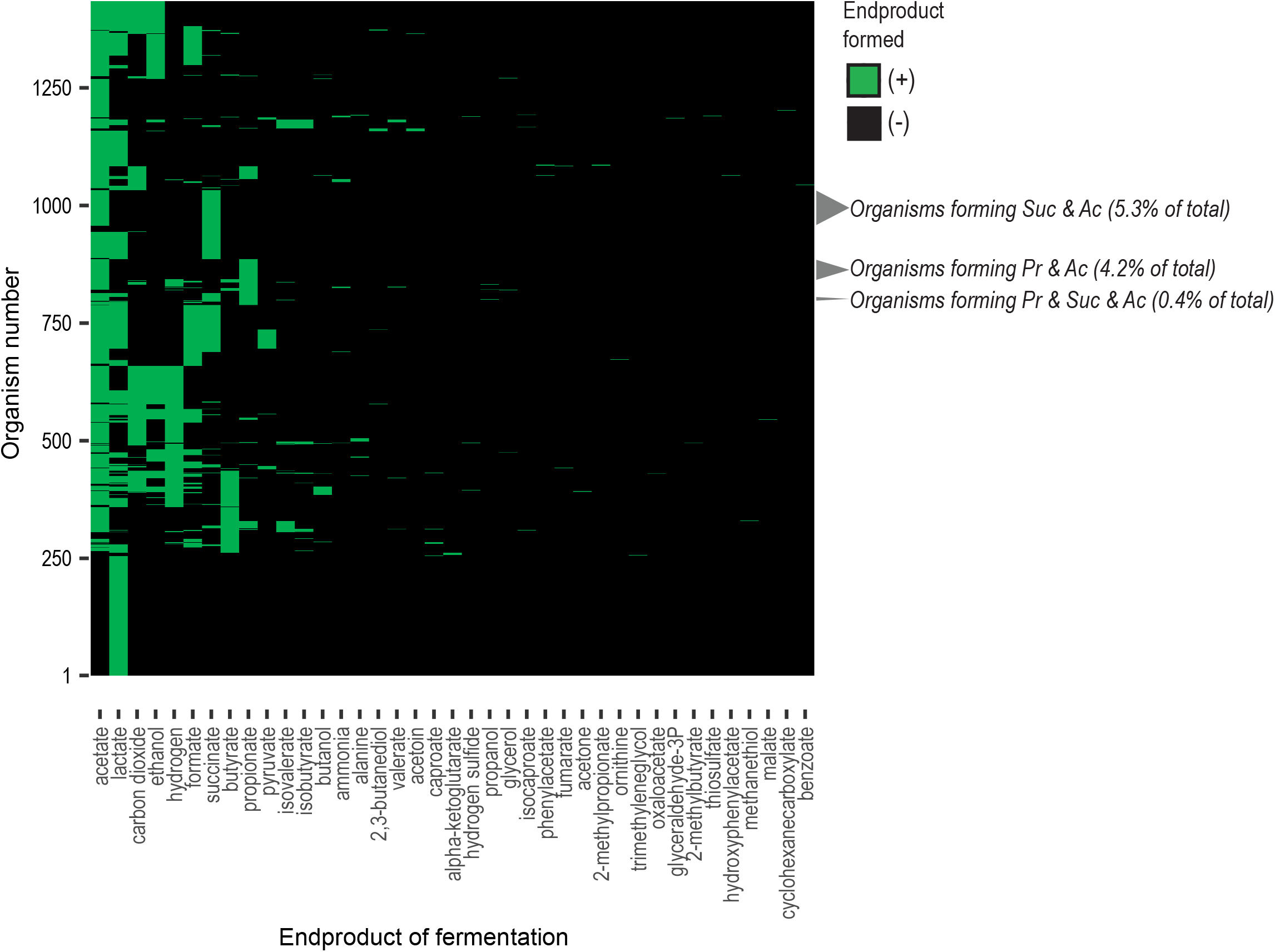
Many organisms form propionate, succinate, and acetate during fermentation. Organisms (n = 1,436) and their reported end products (n = 39) are from *Bergey’s Manual of Systematics of Archaea and Bacteria* (54). Minor (trace) end products are not included. Abbreviations: Ac, acetate; Suc, succinate; Pr, propionate. See Table S5 for more information.

Next, we determined the occurrence of Rnf genes in prokaryotes (Fig. 7, Table S6). We used a total of n = 3,775 type strains for which a genome sequence was available. We found that Rnf genes were uncommon in prokaryotes in general (Fig. 7A). However, these genes were more common in prokaryotes that are fermentative and even more so in those that form propionate, succinate, and acetate. This shows a clear importance of Rnf in such organisms.

**Fig. 7.**
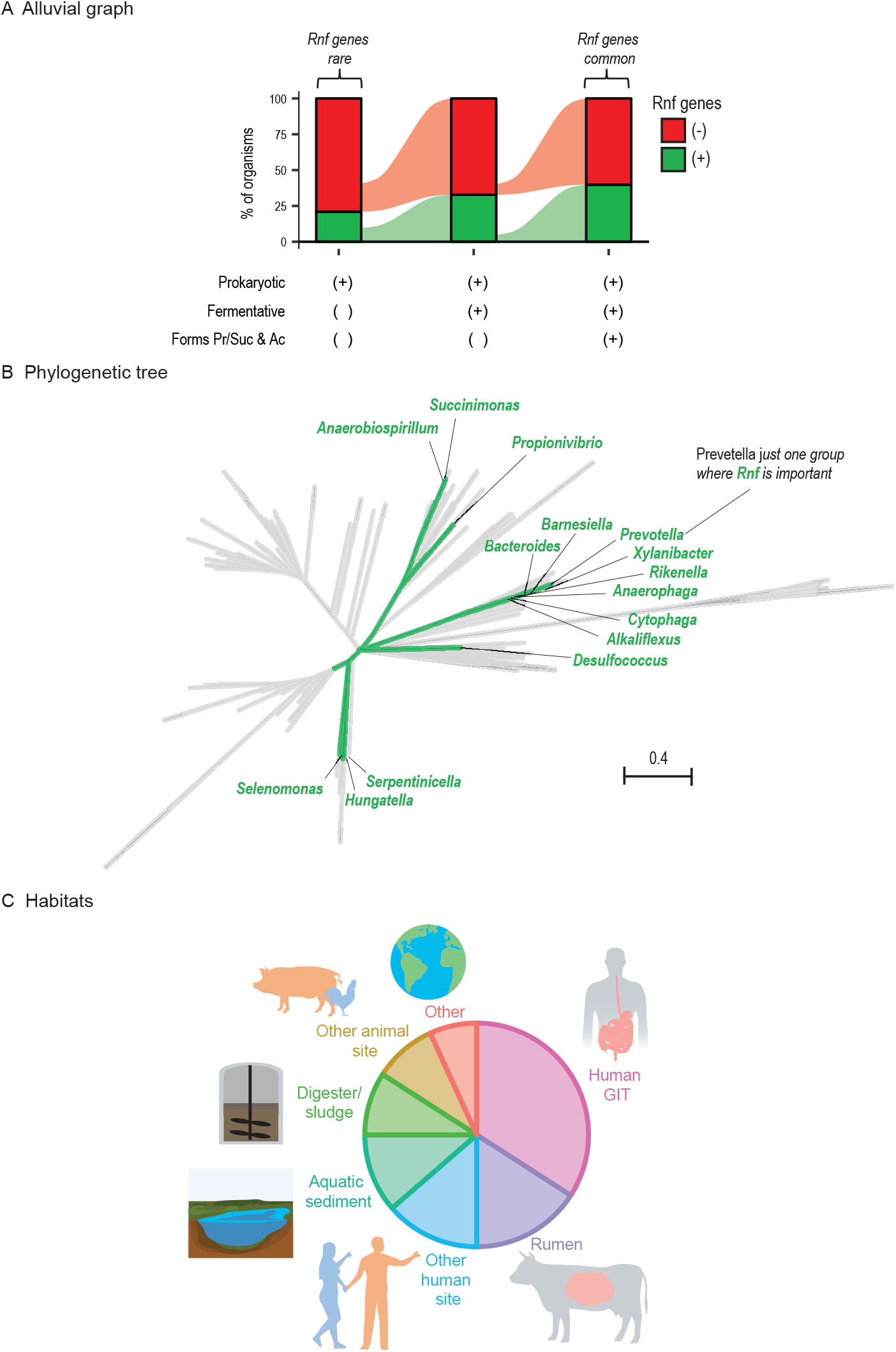
Rnf may be used by many prokaryotes that form propionate, succinate, and acetate during fermentation. (A) Alluvial graph showing percentage of prokaryotes with Rnf genes. Rnf genes are enriched in organisms that are fermentative and form propionate, succinate, and acetate. (B) Phylogenetic tree of prokaryotes, highlighting those with Rnf genes and that form propionate, succinate, and acetate during fermentation. (C) Habitats of prokaryotes with Rnf genes and observed to form propionate, succinate, and acetate during fermentation. Abbreviations: Ac, acetate; Suc, succinate; Pr, propionate. See Tables S5, S6, and S7 for more information.

In total, 44 type strains encoded Rnf and formed propionate, succinate, and acetate during fermentation. A phylogenetic tree shows that these strains are diverse, and they belong to 15 genera (Fig. 7B). Examining their habitats shows that they come from the gut, aquatic sediment, anaerobic digesters, and elsewhere (Fig. 7C, Table S7). Together, these results suggest Rnf is important to propionate formation not just in *Prevotella*, but to many organisms from various habitats.

### Organisms have alternatives to Rnf, but they are uncommon

By oxidizing Fd_red_ and reducing NAD_ox_, Rnf is one enzyme that fills in the missing step of the pathway we study. However, other alternatives can be imagined. To see if any alternatives were common, we used the same genomic and phenotypic data for prokaryotes as before.

We considered five possible pathways (Fig. S3, Table S6, Table S7). One pathway involves the enzyme pyruvate dehydrogenase (EC 1.2.4.1) (Fig. S3A). When this enzyme replaces pyruvate:ferredoxin oxidoreductase, the resulting pathway is balanced without Rnf. This in fact is the pathway originally proposed for propionate formation 60 years ago (32). However, it is uncommon; only 17% of organisms that form propionate, succinate, and acetate encode this enzyme. The four other pathways—involving prototypical hydrogenase (EC 1.12.7.2), bifurcating hydrogenase (EC 1.12.1.4), formate dehydrogenase (EC 1.17.5.3), or *Campylobacter*-type Nuo (33, 34)—are even less common (Fig. S3).

None of these five pathways was found in *Prevotella*. Their genomes did not encode the appropriate enzymes (Tables S3, S4, and S6). We tested cell extracts for catalytic activity of pyruvate dehydrogenase, and we did not find any in *P. brevis* GA33 or *P. ruminicola* 23. We found similar results for cytoplasmic contents and that there was no detectable activity of pyruvate dehydrogenase. As a control, we tested cell extracts of a bacterium with pyruvate dehydrogenase [*E. coli* BL21(DE3)pLysS], and we found high activity (>1 U/mg protein). We also found high activity when spiking cell extract of this bacterium into cell extracts of *P. brevis* GA33 and *P. ruminicola* 23. These controls show our assay worked. Further, we found *P. brevis* GA33 and *P. ruminicola* 23 formed only trace quantities of formate, and they formed no H_2_ (Fig. 2, Fig. S2). In sum, there are no obvious alternatives to Rnf to *Prevotella*. This underscores the importance of Rnf in organisms forming propionate, succinate, and acetate.

## DISCUSSION

Our study shows Rnf is important to forming propionate during fermentation. In *Prevotella*, we show that fermentation is apparently unbalanced and produces excess Fd_red_ and NAD_ox_. Rnf handles the excess Fd_red_ and NAD_ox_ by converting them back to Fd_ox_ and NAD_red_. No other enzyme (or combination of enzymes) had this activity. Rnf thus completes the pathway and allows fermentation to continue.

The pathway for forming propionate has been studied for over 60 years (32), yet the need for an enzyme like Rnf was only recently recognized (17, 18). A likely reason Rnf has been overlooked is the pathway was first elucidated in propionibacteria (32). Propionibacteria have pyruvate dehydrogenase, which, if used, would make the pathway balanced without Rnf (Fig. S3) (32). A pathway with pyruvate dehydrogenase, though plausible, appears seldom used. First, our work shows few organisms forming propionate also encode pyruvate dehydrogenase. Second, propionibacteria themselves may not use this enzyme. Recent work shows they also have pyruvate:ferredoxin oxidoreductase, which is expressed (18, 35) and required for normal growth (18). In *Prevotella*, we found activity of pyruvate:ferredoxin oxidoreductase, but not pyruvate dehydrogenase. Thus, there is a real need for an enzyme like Rnf.

Given this need, we looked for direct evidence of Rnf in *Prevotella*, and we used multiple approaches. Growth experiments showed that fermentation would be unbalanced without Rnf. They showed the problem was serious, and without Rnf, fermentation would halt within 1.5 s. Genomics and proteomics showed that *Prevotella* both encoded and expressed Rnf. Enzyme assays showed *Prevotella* had catalytic activity of Rnf, and its properties were as expected. Our experiments also ruled out alternatives to Rnf. For example, our enzyme assays ruled out presence of a similar enzyme in the cytoplasm [a cytoplasmic ferredoxin--NAD(+) oxidoreductase]. Our growth experiments ruled out H_2_ or formate as other ways of balancing fermentation. In sum, multiple lines of evidence show that *Prevotella* have Rnf and that it balances fermentation.

Our work shows Rnf is encoded by many organisms that form propionate, succinate, and acetate. This result suggests Rnf is important not just to *Prevotella* but many other organisms. Our results also show additional strategies must exist for balancing redox cofactors. We show five strategies, though none is as common as Rnf. Further, we do not examine eukaryotes, even though they have the same problem (19).

Propionate is most commonly formed by succinate pathway, but the acrylate pathway is an alternative. Rnf would be important to either pathway. Though the two pathways involve different carbon intermediates, both produce excess NAD_ox_ and Fd_red_ (2 each per 3 glucose) [see (17)]. Indeed, Rnf is in the proteome of one bacterium that uses the acrylate pathway (36). During ethanol metabolism via the acrylate pathway in a propionate-producer *Anaerotignum neopropionicum*, Rnf is predicted to operate in the reverse direction to reduce Fd_ox_ and oxidize NAD_red_ at the expense of ATP (37). Furthermore, Rnf could be important for NADH regeneration in strains of *Clostridium saccharoperbutylacetonicum* metabolically engineered to produce propionate via the acrylate pathway (38). As an aside, one study suggested that *P. ruminicola* 23 uses the acrylate pathway, not the succinate pathway (39). Our study and others (31) do not support this idea, but Rnf would be important regardless.

The knowledge that Rnf is involved in propionate production is critical for manipulating fermentative propionate production. Modification of metabolic pathways involving redox reactions for synthesis of target metabolites often introduces redox imbalance, which affects the growth and production of the engineered microbes (40). Several cofactor-engineering strategies have been developed to solve the problematic redox imbalance issue (41, 42); however, it is difficult to address this when the knowledge about enzymes involved in redox balance are unknown.

In sum, Rnf completes the pathway for forming propionate formation during fermentation. It has importance in the bacteria we study in the rumen and for bacteria from many other habitats. This work is key to understanding how propionate is formed in the environment and to manipulating its production.

## MATERIALS AND METHODS

### Organisms

*P. brevis* GA33 and *P. ruminicola* 23 were obtained from the ATCC. *Clostridium pasteurianum* 5 and *Pseudobutyrivibrio ruminis* A12-1 were obtained from the DSMZ. *Selenomonas ruminantium* HD4 was obtained from Michael Flythe (USDA-ARS, Lexington, KY) and originally isolated by Marvin Bryant (43). *Escherichia coli* BL21(DE3)pLysS was from Promega.

### Media and growth

Except where noted, strains were grown anaerobically under O_2_-free CO_2_ and with serum bottles with butyl rubber stoppers (44, 45). The inoculant (seed) was 0.1 mL volume of a stationary-phase culture. The temperature of growth was 37°C.

*P. brevis* GA33 and *S. ruminantium* HD4 were cultured on the medium PC+VFA (46). *P. ruminis* A12-1 was cultured on a complex medium as described by Schoelmerich et al. (27). *P. ruminicola* 23 was cultured on medium BZ. We developed this defined medium from a complex medium (47). Per liter, the medium contained 8 g glucose, 0.6 g K_2_HPO_4_, 0.45 g KH_2_PO_4_, 0.45 g (NH_4_)_2_SO_4_, 0.9 g NaCl, 92 mg MgSO_4_, 0.12 g CaCl_2_·2H_2_O, 2 mL of 0.5 g/L hemin in 10 mM NaOH, 1 mL 0.1% (w/v) resazurin, 1 mL trace element SL-9 (48), 10 mL DSMZ-medium-141 Wolin’s vitamin solution, 0.1 mg vitamin B_12_, 322.7 μL isobutyric acid, 322.7 μL 2-methylbutyric acid, 322.7 μL valeric acid, 322.7 μL isovaleric acid, 4 g Na_2_CO_3_, and 1.2 g L-cysteine·HCl·H_2_O. Glucose, Wolin’s vitamin solution, and vitamin B_12_ were added to medium BZ after autoclaving. *C. pasteurianum* 5 was cultured on a glucose medium in 1-L Pyrex bottle sealed with stoppers. Per liter, the medium contained 20 g glucose, 15.329 g K_2_HPO_4_, 1.5 g KH_2_PO_4_, 0.1 g NaCl, 98 mg MgSO_4_, 10 mg Na_2_MoO_4_·2H_2_O, 1 g NH_4_Cl, 50 mg FeSO_4_·7 H_2_O, 5 mg 4-aminobenzoic acid, and 1 mg biotin. *E. coli* BL21(DE3)pLysS was cultured aerobically on Luria-Bertani medium.

Growth of cultures was measured by removing 1-mL aliquots with a syringe and measuring optical density at 600 nm (OD_600_) in cuvettes in a Thermo Scientific Genesys 20 spectrophotometer. The sample was diluted with 0.9% (w/v) NaCl as needed to remain within the linear range of the instrument.

### Analysis of fermentation products and cells

Three, 70-mL cultures were inoculated and grown to the late-log phase (OD_600_ = 1.3 for *P. brevis* GA33 and OD_600_ = 4.0 for *P. ruminicola* 23). Cells were harvested by centrifugation (21,100 × *g* for 20 min at 4°C). The supernatant was stored at -20°C. Cell pellets were resuspended in ddH_2_O and harvested by centrifugation (21,100 × *g* for 30 min at 4°C). Pellets were transferred to aluminum pans with ddH_2_O and dried at 105 °C overnight. The dry mass of cells was determined by weighing the pan with dried pellet (while still hot) (49). After cooling, an aliquot of pellet was submitted for elemental analysis (C, H, N) by Intertek (Whitehouse, NJ). The cooled pellet was reweighed to correct for any water absorbed.

Supernatant was analyzed for glucose and fermentation products according to Zhang et al. (50) with modifications. Specifically, acetate was measured by gas chromatography rather than enzymatic assay. Ethanol was measured with a commercial kit from Megazyme (product code K-ETOH).

One aliquot of culture (5-mL) was also collected at the start of the incubation. Cells were removed, and supernatant was analyzed as above. The inoculant for cultures was 0.1 mL of a late-log phase culture. The dry mass of cells in this inoculant was determined by methods above. The elemental composition (C, H, N) was assumed to be the same as cells inoculated and grown to the late-log phase.

### Recovery of carbon and hydrogen

We calculated recovery of carbon in cells and fermentation products. Recovery is defined as the (total carbon at end)/(total carbon at start) × 100%.

Total carbon (mmol C L^-1^) was the sum of carbon in cells, glucose, fermentation acids, and CO_2_. For CO_2_, we defined the concentration at the start as 0. The concentration at the end was calculated from stoichiometry, assuming -1 CO_2_/formate, 1 CO_2_/acetate, -1 CO_2_/succinate, 2 CO_2_/butyrate, 2 CO_2_/isobutyrate, 1 CO_2_/valerate, 1 CO_2_/isovalerate, and 1 CO_2_/ethanol [after Hackmann et al. (51)]. CO_2_ formed during cell synthesis was ignored.

Recovery of hydrogen was calculated analogously. For H_2_O, we defined the concentration at the start (mmol H L^-1^) as 0. We calculated the concentration at the end (mmol H L^-1^) from stoichiometry, assuming 1 H_2_O/acetate, 1 H_2_O/propionate, and 1 H_2_O/succinate [after Hackmann et al. (51)]. H_2_O formed during cell synthesis was ignored.

### Rate of glucose fermentation

We measured rate of glucose fermentation by *P. ruminicola* 23 in the mid-exponential phase. Samples of culture were collected at 4 points during this phase [where ln(OD_600_) increased linearly over time]. The glucose concentration (mmol L^-1^) was measured as above. The dry cell weight (g dry cell L^-1^) was calculated from OD_600_ (referring to samples where both OD_600_ and weight were known). The rate of glucose consumption [nmol glucose (g dry cell)^-1^ s^-1^] was directly calculated. The rate of glucose fermentation was assumed to be glucose consumption × 0.642 (see Table S1). The final value for three biological replicates was 667 nmol glucose fermented (g dry cell)^-1^ s^-1^.

### Proteomics

We used proteomics to determine what genes were expressed in the cells of *P. brevis* GA33 and *P. ruminicola* 23. Peptide samples from cell extract and cell membrane were prepared and analyzed using LC-MS (see Text S1 for details).

### Enzyme assays

We measured activities of enzymes in cell extract, cell membrane, and cytoplasmic contents. Assays were performed following Zhang et al (50). The temperature and other conditions were as reported in Table 1. One unit of activity is defined as 1 μmol of product formed per min. To correct Rnf activity for NADH dehydrogenase activity, we added to its value the activity measured for Nqr (see Table 2).

**TABLE 1.**
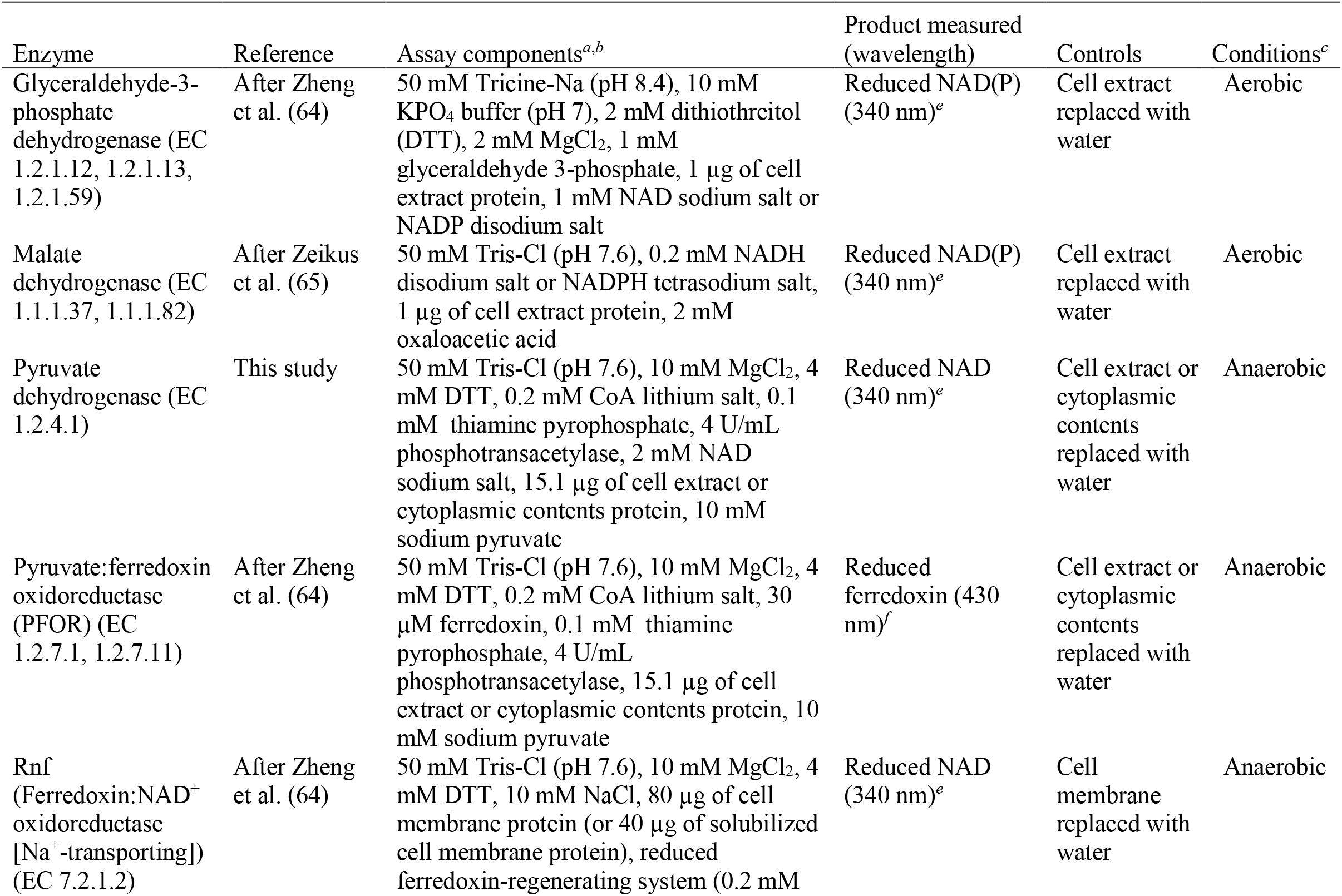
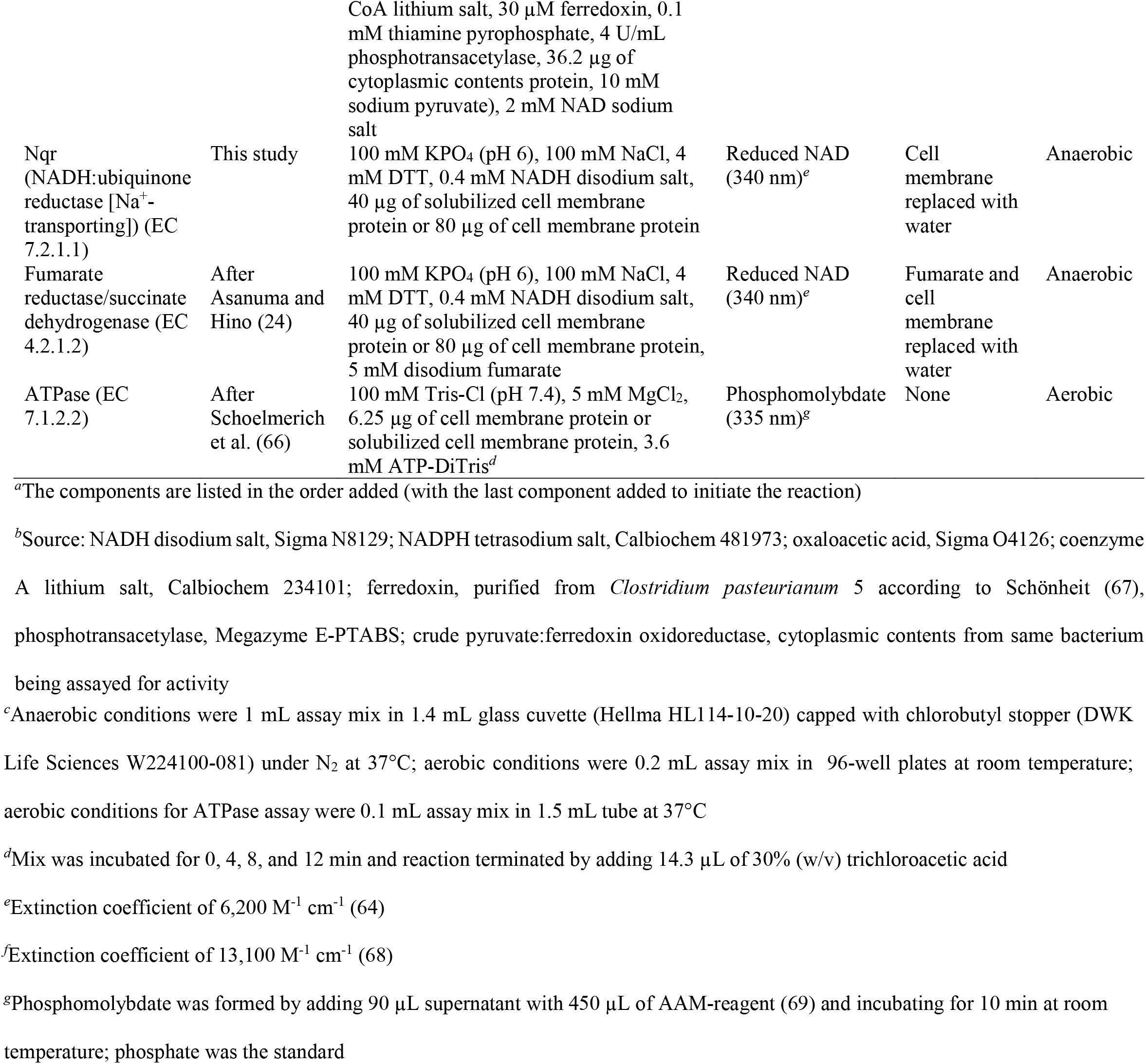
Conditions used to measure enzymatic activity

| Enzyme | Reference | Assay components<sup>a,b</sup> | Product measured (wavelength) | Controls | Conditions<sup>c</sup> |
| --- | --- | --- | --- | --- | --- |
| Glyceraldehyde-3-phosphate dehydrogenase (EC 1.2.1.12, 1.2.1.13, 1.2.1.59) | After Zheng et al. (64) | 50 mM Tricine-Na (pH 8.4), 10 mM KPO<sub>4</sub> buffer (pH 7), 2 mM dithiothreitol (DTT), 2 mM MgCl<sub>2</sub>, 1 mM glyceraldehyde 3-phosphate, 1 µg of cell extract protein, 1 mM NAD sodium salt or NADP disodium salt | Reduced NAD(P) (340 nm)<sup>e</sup> | Cell extract replaced with water | Aerobic |
| Malate dehydrogenase (EC 1.1.1.37, 1.1.1.82) | After Zeikus et al. (65) | 50 mM Tris-Cl (pH 7.6), 0.2 mM NADH disodium salt or NADPH tetrasodium salt, 1 µg of cell extract protein, 2 mM oxaloacetic acid | Reduced NAD(P) (340 nm)<sup>e</sup> | Cell extract replaced with water | Aerobic |
| Pyruvate dehydrogenase (EC 1.2.4.1) | This study | 50 mM Tris-Cl (pH 7.6), 10 mM MgCl<sub>2</sub>, 4 mM DTT, 0.2 mM CoA lithium salt, 0.1 mM thiamine pyrophosphate, 4 U/mL phosphotransacetylase, 2 mM NAD sodium salt, 15.1 µg of cell extract or cytoplasmic contents protein, 10 mM sodium pyruvate | Reduced NAD (340 nm)<sup>e</sup> | Cell extract or cytoplasmic contents replaced with water | Anaerobic |
| Pyruvate:ferredoxin oxidoreductase (PFOR) (EC 1.2.7.1, 1.2.7.11) | After Zheng et al. (64) | 50 mM Tris-Cl (pH 7.6), 10 mM MgCl<sub>2</sub>, 4 mM DTT, 0.2 mM CoA lithium salt, 30 µM ferredoxin, 0.1 mM thiamine pyrophosphate, 4 U/mL phosphotransacetylase, 15.1 µg of cell extract or cytoplasmic contents protein, 10 mM sodium pyruvate | Reduced ferredoxin (430 nm)<sup>f</sup> | Cell extract or cytoplasmic contents replaced with water | Anaerobic |
| Rnf (Ferredoxin:NAD<sup>+</sup> oxidoreductase [Na<sup>+</sup>-transporting]) (EC 7.2.1.2) | After Zheng et al. (64) | 50 mM Tris-Cl (pH 7.6), 10 mM MgCl<sub>2</sub>, 4 mM DTT, 10 mM NaCl, 80 µg of cell membrane protein (or 40 µg of solubilized cell membrane protein), reduced ferredoxin-regenerating system (0.2 mM | Reduced NAD (340 nm)<sup>e</sup> | Cell membrane replaced with water | Anaerobic |
| | | CoA lithium salt, 30 μM ferredoxin, 0.1 mM thiamine pyrophosphate, 4 U/mL phosphotransacetylase, 36.2 μg of cytoplasmic contents protein, 10 mM sodium pyruvate), 2 mM NAD sodium salt | | | |
| Nqr (NADH:ubiquinone reductase [$\text{Na}^+$-transporting]) (EC 7.2.1.1) | This study | 100 mM $\text{KPO}_4$ (pH 6), 100 mM NaCl, 4 mM DTT, 0.4 mM NADH disodium salt, 40 μg of solubilized cell membrane protein or 80 μg of cell membrane protein | Reduced NAD (340 nm)<sup>e</sup> | Cell membrane replaced with water | Anaerobic |
| Fumarate reductase/succinate dehydrogenase (EC 4.2.1.2) | After Asanuma and Hino (24) | 100 mM $\text{KPO}_4$ (pH 6), 100 mM NaCl, 4 mM DTT, 0.4 mM NADH disodium salt, 40 μg of solubilized cell membrane protein or 80 μg of cell membrane protein, 5 mM disodium fumarate | Reduced NAD (340 nm)<sup>e</sup> | Fumarate and cell membrane replaced with water | Anaerobic |
| ATPase (EC 7.1.2.2) | After Schoelmerich et al. (66) | 100 mM Tris-Cl (pH 7.4), 5 mM $\text{MgCl}_2$, 6.25 μg of cell membrane protein or solubilized cell membrane protein, 3.6 mM ATP-DiTris<sup>d</sup> | Phosphomolybdate (335 nm)<sup>g</sup> | None | Aerobic |

<sup>a</sup>The components are listed in the order added (with the last component added to initiate the reaction)
<sup>b</sup>Source: NADH disodium salt, Sigma N8129; NADPH tetrasodium salt, Calbiochem 481973; oxaloacetic acid, Sigma O4126; coenzyme A lithium salt, Calbiochem 234101; ferredoxin, purified from *Clostridium pasteurianum* 5 according to Schönheit (67), phosphotransacetylase, Megazyme E-PTABS; crude pyruvate:ferredoxin oxidoreductase, cytoplasmic contents from same bacterium being assayed for activity
<sup>c</sup>Anaerobic conditions were 1 mL assay mix in 1.4 mL glass cuvette (Hellma HL114-10-20) capped with chlorobutyl stopper (DWK Life Sciences W224100-081) under N<sub>2</sub> at 37°C; aerobic conditions were 0.2 mL assay mix in 96-well plates at room temperature; aerobic conditions for ATPase assay were 0.1 mL assay mix in 1.5 mL tube at 37°C <sup>d</sup>Mix was incubated for 0, 4, 8, and 12 min and reaction terminated by adding 14.3 µL of 30% (w/v) trichloroacetic acid <sup>e</sup>Extinction coefficient of 6,200 M<sup>-1</sup> cm<sup>-1</sup> (64) <sup>f</sup>Extinction coefficient of 13,100 M<sup>-1</sup> cm<sup>-1</sup> (68) <sup>g</sup>Phosphomolybdate was formed by adding 90 µL supernatant with 450 µL of AAM-reagent (69) and incubating for 10 min at room temperature; phosphate was the standard

**TABLE 2.**
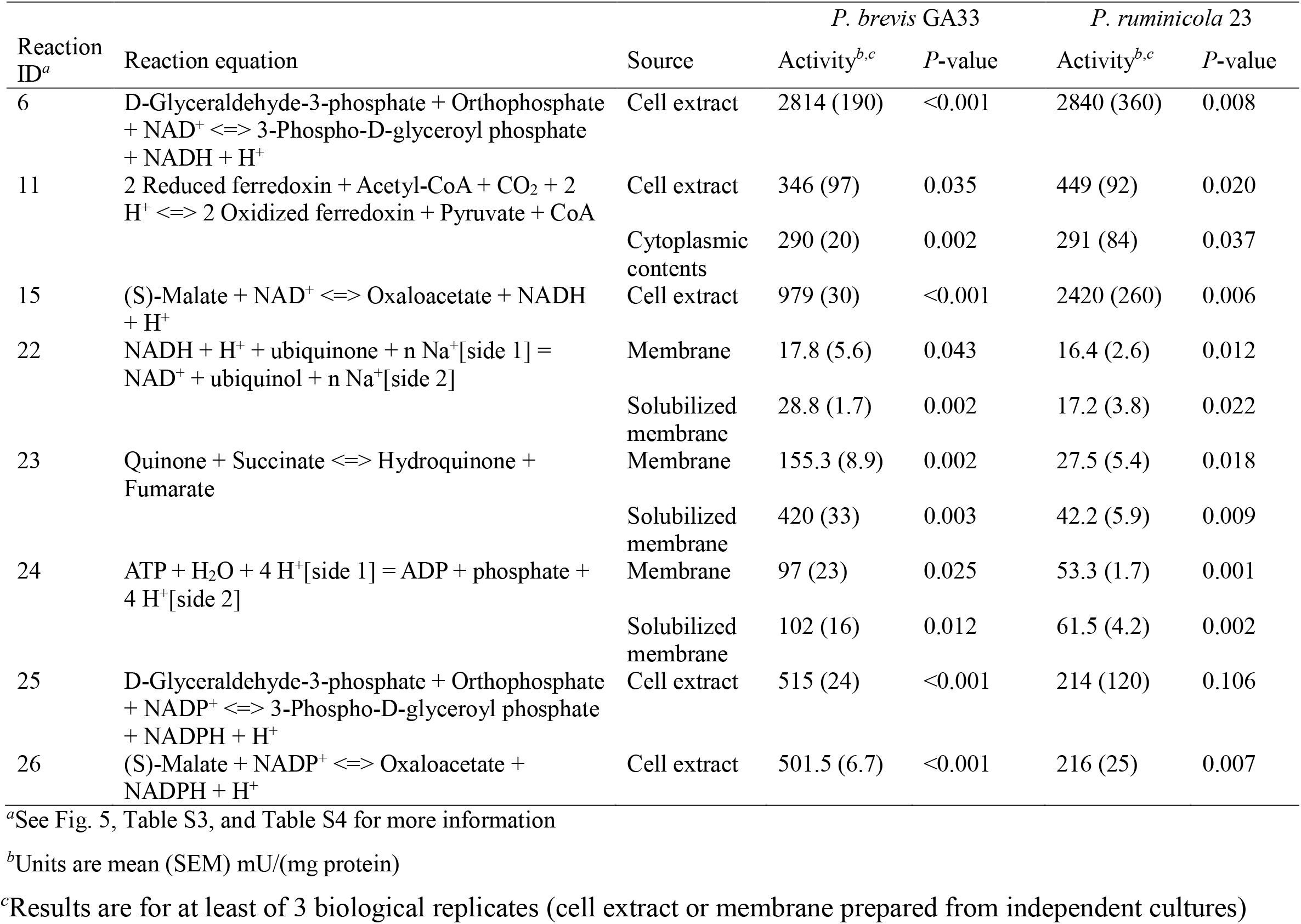
Enzymatic assays confirm *Prevotella* catalyze key reactions for forming propionate, succinate, and acetate

| Reaction ID<sup>a</sup> | Reaction equation | Source | *P. brevis* GA33 |  | *P. ruminicola* 23 |  |
| --- | --- | --- | --- | --- | --- | --- |
|  |  |  | Activity<sup>b,c</sup> | P-value | Activity<sup>b,c</sup> | P-value |
| 6 | D-Glyceraldehyde-3-phosphate + Orthophosphate + NAD<sup>+</sup> <=> 3-Phospho-D-glyceroyl phosphate + NADH + H<sup>+</sup> | Cell extract | 2814 (190) | <0.001 | 2840 (360) | 0.008 |
| 11 | 2 Reduced ferredoxin + Acetyl-CoA + CO<sub>2</sub> + 2 H<sup>+</sup> <=> 2 Oxidized ferredoxin + Pyruvate + CoA | Cell extract | 346 (97) | 0.035 | 449 (92) | 0.020 |
|  |  | Cytoplasmic contents | 290 (20) | 0.002 | 291 (84) | 0.037 |
| 15 | (S)-Malate + NAD<sup>+</sup> <=> Oxaloacetate + NADH + H<sup>+</sup> | Cell extract | 979 (30) | <0.001 | 2420 (260) | 0.006 |
| 22 | NADH + H<sup>+</sup> + ubiquinone + n Na<sup>+</sup>[side 1] = NAD<sup>+</sup> + ubiquinol + n Na<sup>+</sup>[side 2] | Membrane | 17.8 (5.6) | 0.043 | 16.4 (2.6) | 0.012 |
|  |  | Solubilized membrane | 28.8 (1.7) | 0.002 | 17.2 (3.8) | 0.022 |
| 23 | Quinone + Succinate <=> Hydroquinone + Fumarate | Membrane | 155.3 (8.9) | 0.002 | 27.5 (5.4) | 0.018 |
|  |  | Solubilized membrane | 420 (33) | 0.003 | 42.2 (5.9) | 0.009 |
| 24 | ATP + H<sub>2</sub>O + 4 H<sup>+</sup>[side 1] = ADP + phosphate + 4 H<sup>+</sup>[side 2] | Membrane | 97 (23) | 0.025 | 53.3 (1.7) | 0.001 |
|  |  | Solubilized membrane | 102 (16) | 0.012 | 61.5 (4.2) | 0.002 |
| 25 | D-Glyceraldehyde-3-phosphate + Orthophosphate + NADP<sup>+</sup> <=> 3-Phospho-D-glyceroyl phosphate + NADPH + H<sup>+</sup> | Cell extract | 515 (24) | <0.001 | 214 (120) | 0.106 |
| 26 | (S)-Malate + NADP<sup>+</sup> <=> Oxaloacetate + NADPH + H<sup>+</sup> | Cell extract | 501.5 (6.7) | <0.001 | 216 (25) | 0.007 |

<sup>a</sup>See Fig. 5, Table S3, and Table S4 for more information
<sup>b</sup>Units are mean (SEM) mU/(mg protein)
<sup>c</sup>Results are for at least of 3 biological replicates (cell extract or membrane prepared from independent cultures)

Samples (cell extract, cell membrane, cytoplasmic contents) were prepared according to Text S1. When required, ferredoxin was purified from *C. pasteurianum* 5 was according to reference (52) with modifications (see Text S1).

### Other chemical analyses

Protein was measured using the Bradford method (53). The standard was bovine serum albumin. H_2_ was measured with gas chromatography (see Text S1).

### Information for organisms in Bergey’s Manual

We collected phenotypic, genomic, and other information for organisms in *Bergey’s Manual of Systematics of Archaea and Bacteria* (54). All n = 1,836 articles for genera in *Bergey’s Manual* was downloaded. Names and written descriptions of n = 8,026 type strains were then extracted from the text. We used R scripts from Hackmann and Zhang (55) to automate this process. To collect phenotypic information, we read written descriptions of the type strains. This information included fermentative ability and major fermentation endproducts. To collect genomic information, R scripts from Hackmann and Zhang (55) were used. The scripts first extracted out the organism’s taxonomy and article link from the written description. The scripts then used the taxonomy to find an organism’s GOLD organism ID, GOLD project ID, and IMG genome ID.

### Searches for genes and proteins

We searched genomes for genes involved in forming propionate, succinate, and acetate. To do so, we used IMG/M database (56), the IMG/M genome ID for each genome, and the KEGG Orthology (KO) ID for each gene (57). For some genes, we searched for the COG (58) or pfam (59) ID instead. For hydrogenases, we followed methods in Text S1.

For each gene, we report the respective enzyme name, enzyme symbol, EC number, and biochemical reaction. This information came from KEGG (57) and HydDB (60). An enzyme was considered present in the genome if genes for all subunits was found. A reaction was considered present if at least one isozyme was found.

### Other bioinformatic analyses

Proteomes were searched for proteins using locus tags for genes above. Phylogenetic trees were constructed according to Hackmann and Zhang (55). We identified habitats of organisms forming propionate, succinate, and acetate using *Bergey’s Manual* (54), BacDive (61), and information from public culture collections. Structures of proteins were predicted using ColabFold (62), then they were visualized with PyMOL according to Hackmann (63).

### Statistics

A one-sided *t*-tests was used to determine if mean yield of fermentation products and mean values of enzymatic activity was greater than 0. *P*-values reported are for that test.

### Data availability

The LC-MS data have been deposited in the Proteomics Identification (PRIDE) Archive with the dataset identifier PXD034119.

## ACKNOWLEDGMENTS

We thank Dr. Gabriela Grigorean of the UC Davis Proteomics Core for performing LC-MS analysis. This work was supported by an Agriculture and Food Research Initiative Competitive Grant [grant no. 2018-67015-27495] and Hatch Project [accession no. 1019985] from the United States Department of Agriculture National Institute of Food and Agriculture.

