## Supplementary Figures for "The enzyme Rnf completes the pathway for forming propionate during fermentation in *Prevotella*"

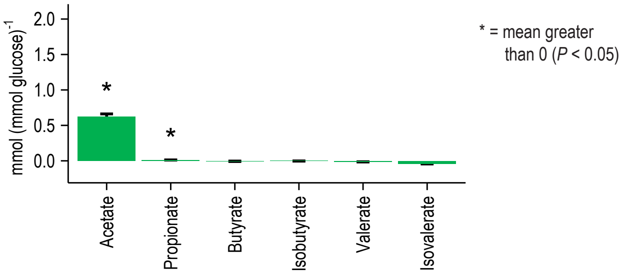


**Fig. S1.** *Prevotella brevis* GA33 forms trace amounts of propionate. Experiments are as in Fig. 2, except acetate and propionate were withheld from the medium PC+VFA. Propionate is in high concentrations in normal media (c. 6.4 mmol $L^{-1}$), and withholding it allowed propionate formation to be detected more sensitively. Products not shown were not measured.

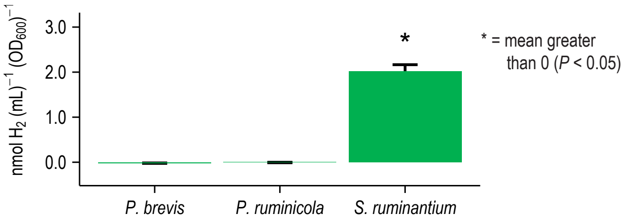


**Fig. S2.** *Prevotella* do not form $H_2$ during fermentation of glucose. *Selenomonas ruminantium* HD4 is known to form trace amounts of $H_2$ (Scheifinger et al 1975) and is included as a control.

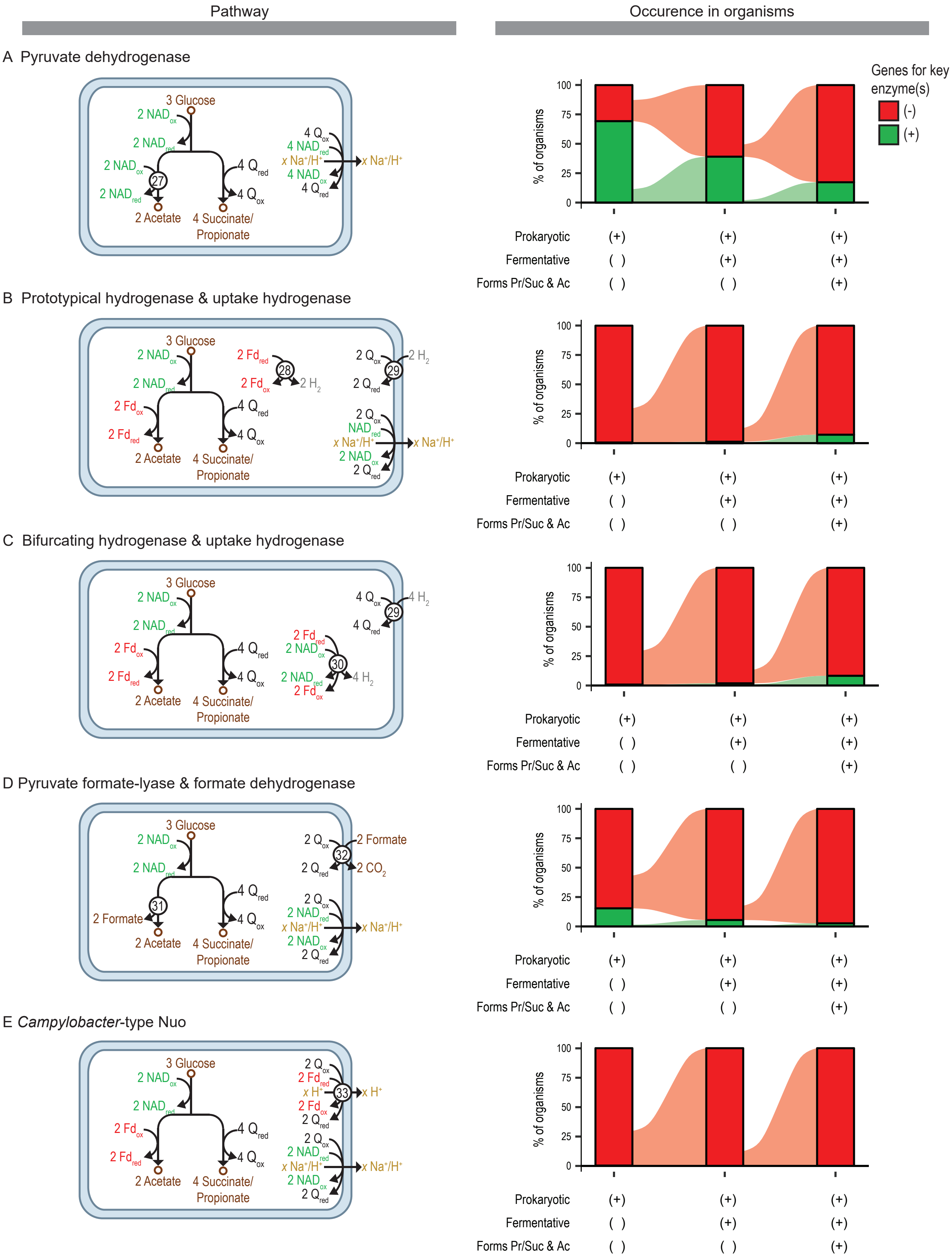


**Fig. S3.** Rnf is not needed in all pathways that form propionate, succinate, and acetate, but these alternatives are uncommon. Alternatives to Rnf involve (A) pyruvate dehydrogenase, (B) prototypical hydrogenase and uptake hydrogenase, (C) bifurcating hydrogenase and uptake hydrogenase, (D) pyruvate formate-lyase and formate dehydrogenase, and (E) *Campylobacter*-type Nuo. Conversion of $Q_{red}$ to $Q_{ox}$ is drawn in middle of cell, but it actually occurs at membrane. Abbreviations: $Fd_{ox}$, oxidized ferredoxin; $Fd_{red}$, reduced ferredoxin (two reduced iron-sulfur clusters); $NAD_{ox}$, oxidized NAD; $NAD_{red}$, reduced NAD; $Q_{ox}$, oxidized quinone; $Q_{red}$, reduced quinone.
